# Intramuscular adipose tissue physically restricts functional muscle recovery

**DOI:** 10.1101/2024.12.17.628009

**Authors:** A.M. Norris, V.R. Palzkill, A.B. Appu, T.E. Ryan, D. Kopinke

## Abstract

With age and disease, skeletal muscle is progressively lost and replaced by fibrotic scar and intramuscular adipose tissue (IMAT). While strongly correlated, it remains unclear whether IMAT has a functional impact on muscle. In the present study, we evaluated the effects of IMAT during muscle injury by creating a mouse model where the cellular origin of IMAT, fibro/adipogenic progenitors (FAPs), are prevented from differentiating into adipocytes (FATBLOCK model). We found that blocking IMAT after an adipogenic injury allowed muscle to regenerate more efficiently, resulting in enhanced function. Our data explain why acute muscle injuries featuring IMAT infiltration, such as rotator cuff tears and acute denervation injuries, exhibit poor regeneration and lead to a loss in muscle function. It also demonstrates the therapeutic importance of preventing IMAT formation in acute injuries in order to maximize regeneration and minimize loss in muscle mass and function.

## INTRODUCTION

Adipose tissue is an important organ, acting as an energy storage, as well as an endocrine organ, secreting hormones that maintain a healthy organism (reviewed in (Kershaw and Flier, 2004)). However, different fat depots, such as subcutaneous (SAT), visceral (VAT) and intramuscular fat (IMAT) have stark metabolic and phenotypic differences (Bjørndal et al., 2011; Ford et al., 2023; Sachs et al., 2019). IMAT, the accumulation of adipocytes between individual myofibers of skeletal muscle, is a pathological hallmark of muscular dystrophies (Hogarth et al., 2019; Milad et al., 2017; Willcocks et al., 2016; Wokke et al., 2014), but is also present in a spectrum of metabolic disorders including diabetes and obesity, and sarcopenia (Goodpaster et al., 2005; Goodpaster et al., 2003; Goodpaster et al., 2006; Goodpaster et al., 2000a; Goodpaster et al., 2000b; Murphy et al., 1986a). The progressive infiltration of IMAT within muscle tissue has been closely associated with loss of muscle mass, metabolic dysfunction, disease progression and impairment of patient mobility (Barnard et al., 2020; Lim et al., 2019; Park et al., 2007). In addition to being a feature of chronic diseases, IMAT also arises in response to acute insults such as spinal cord and rotator cuff injuries, where muscle tissue is also simultaneously lost (Beeler et al., 2013; Gibbons et al., 2018; Thangarajah et al., 2017). Despite the strong correlation between IMAT and loss of muscle tissue, it is not fully understood whether IMAT has both a direct and negative effect on muscle, or whether it is just a neutral bystander, filling the space lost by dying myofibers.

The cellular origin of IMAT is a population of stem cells located in the muscle interstitium, called fibro-adipogenic progenitors (FAPs) (Flores-Opazo et al., 2024; Murphy et al., 2011; Pisani et al., 2010; Uezumi et al., 2014b; Uezumi et al., 2010; Uezumi et al., 2011). In a healthy muscle, FAPs are critical in maintaining muscle mass during homeostasis and play a central role in muscle regeneration (Leinroth et al., 2022; Murphy et al., 2011; Wosczyna et al., 2019). During early regeneration, FAPs secrete pro-myogenic factors to aid the cellular origin of muscle fibers, muscle stem cells (MuSCs), in their differentiation process towards myofibers. With age and disease, however, FAPs can also differentiate into either adipocytes, leading to IMAT formation, or myofibroblasts, giving rise to fibrosis (Flores-Opazo et al., 2024; Murphy et al., 2011; Pisani et al., 2010; Uezumi et al., 2014b; Uezumi et al., 2010; Uezumi et al., 2011). While it is not completely understood what drives FAPs to differentiate into IMAT in certain conditions, the downstream adipogenic signaling cascade is governed by *Pparγ,* the master adipogenic regulator (Barak et al., 1999; Rosen et al., 1999b). *Pparγ* is a transcription factor which, when active, initiates the signaling cascade required for the commitment and subsequent differentiation of adipogenic progenitors towards the adipogenic lineage (Siersbæk et al., 2010). Given the widespread presence of IMAT in pathological conditions, there is a pressing need to further understand the different etiologies in which IMAT arises and what its effects are on skeletal muscle.

In the present study, we created a conditional mouse model, termed FATBLOCK, where IMAT is blocked by preventing FAPs from differentiating into adipocytes through the inducible genetic removal of *Pparγ*. Under homeostatic conditions, this deletion caused no gross effects on overall health, lipid imbalance or muscle function. However, upon an adipogenic injury, deletion of *Pparγ* from FAPs blocks their differentiation into adipocytes in a cell autonomous manner, leading to a significant repression in IMAT formation. When assessing the effects of IMAT on muscle, we found that IMAT acted as a physical barrier and prevented new nascent myofibers from forming during early regeneration, leading to a reduction in myofiber density that persisted at later stages. In addition, IMAT interfered with myofiber hypertrophy post regeneration resulting in smaller myofibers at 21 days post injury. Taken together, we found that in the context of an acute adipogenic injury, IMAT has a direct and negative effect on muscle regeneration by physically preventing myofiber formation during early regeneration, as well as myofiber hypertrophy during the later regenerative phase. Consequently, this results in a functionally weakened muscle that has both fewer and smaller myofibers.

## RESULTS

### Removal of *Pparγ* from FAPs has no detectable impact on muscle or overall health during homeostasis

To determine the functional impact of IMAT on muscle, we created a mouse model where FAPs are prevented from differentiating into adipocytes (Fig. 1A). For this, we crossed the FAP-specific tamoxifen-inducible *Pdgfra^CreERT2^*allele (Chung et al., 2018) to a conditional allele of *Pparγ* (He et al., 2003), the master regulator of adipogenesis (Barak et al., 1999; Rosen et al., 1999b) (*Pdgfra^CreERT^ Pparγ^lox/lox^*, referred to as FAP^no^ ^Pparγ^ or Pparγ^Δ/Δ^). PDGFRA is the “gold-standard” marker in the field to identify FAPs within murine and human muscle (Hogarth et al., 2019; Joe et al., 2010; Kopinke et al., 2017; Lukjanenko et al., 2019; Santini et al., 2020; Uezumi et al., 2014a; Uezumi et al., 2010; Uezumi et al., 2011; Wosczyna et al., 2019). Mice lacking the Cre allele served as controls. Tamoxifen (TMX) was administered by oral gavage on two consecutive days and 4 weeks later, the overall health of the mouse was assessed, as well as its muscle function (Fig 1A). We found no difference in total body weight or tibialis anterior (TA) weight in females or males between wild type (WT) and FAP^no^ ^Pparγ^ mice (Fig. 1B). Thus, short term loss of *Pparγ* from FAPs has no gross effect on overall health or muscle mass.

**Figure 1.**
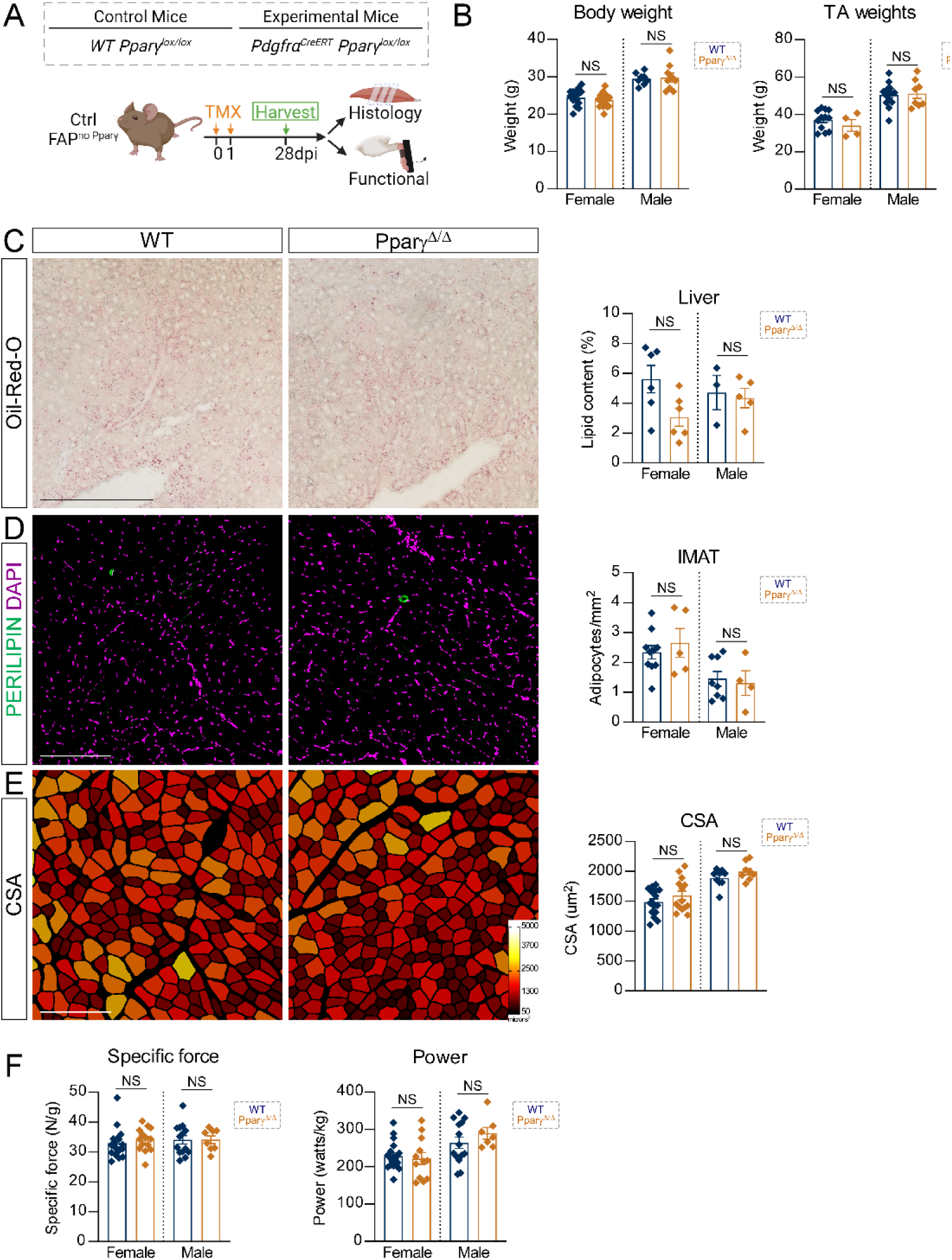
FAP-specific deletion of *Pparγ* has no overall effects during homeostasis. **A)** Experimental design. **B)** (*left*) Whole body weight of experimental mice (g) (females: WT n= 16 mice and _FAPno Pparγ n=16_ mice; males: WT n=8 mice and _FAPno Pparγ n=9_ mice). (*right*) Weight of tibialis anterior (TA) muscle (mg) (females: WT n=12 TAs and FAP^no Pparγ^ n=4 TAs; males: WT n=14 TAs and _FAPno Pparγ n=8_ TAs). Weights recorded 28 days after tamoxifen administration. **C)** (*left*) Cross-section of liver from WT and FAP^no Pparγ^ mice stained with Oil-Red-O; scale bar: 250µm. (*right*) Quantification of lipid content as a percentage of area occupied by Oil-Red-O staining (females: WT n= 6 mice and FAP^no^ ^Pparγ^ n=6 mice; males: WT n=3 mice and _FAPno Pparγ n=5_ mice). **D)** Immuno-fluorescence of cross-sections of TAs stained for adipocytes (PERILIPIN, green) and nuclei visualized through DAPI (magenta). Scale bar: 200µm. Quantification of number of adipocytes normalized to TA area (females: WT n= 10 TAs and FAP^no Pparγ^ n=5 TAs; males: WT n=8 TAs and FAP^no Pparγ^ n=4 TAs). **E)** Myofibers color-coded according to their cross-sectional area in WT and FAP^no Pparγ^ mice. Scale bars: 200µm. Quantification of average cross-sectional area (CSA) (females: WT n=16 TAs and FAP^no Pparγ^ n=14 TAs; males: WT n=10 TAs and FAP^no Pparγ^ n=8 TAs). **F)** Quantification of *in vivo* force production for (left) specific force (females: WT n=17 TAs and FAP^no Pparγ^ n=14 TAs; males: WT n=14 TAs and FAP^no Pparγ^ n=8 TAs) and power (females: WT n=20 TAs and FAP^no Pparγ^ n=12 TAs; males: WT n=14 TAs and FAP^no Pparγ^ n=8 TAs). All data are represented as mean ± SEM. A multiple unpaired two-tailed t test followed by a Holm-Šídák post hoc test was used.

Previous attempts to determine the function of IMAT on muscle focused on a cell ablation approach, which resulted in complete loss of all adipocytes throughout the body (Biltz et al., 2020). Unfortunately, this resulted in systemic lipid imbalance and caused lipids to accumulate in the liver, known as hepatic steatosis. To investigate whether loss of *Pparγ* from FAPs leads to widespread lipid imbalance, we assessed lipid deposition within the liver, visualized through an Oil-Red-O staining, in FAP^no Pparγ^ mice and WT littermates (Fig. 1C). We found no difference in hepatic lipid deposition between genotypes (Fig. 1C), indicating that deletion of *Pparγ* within FAPs does not cause systemic lipid dysregulation.

We next evaluated the impact FAP-specific *Pparγ* deletion has on IMAT formation under homeostatic conditions by quantifying PERILIPIN^+^ adipocytes normalized to total TA area (Fig. 1D). We found no significant difference between genotypes of either sex (Fig. 1D), demonstrating that *Pparγ* deletion in FAPs does not affect IMAT homeostasis or *de novo* adipogenic formation under homeostatic conditions.

Finally, we assessed the effects of *Pparγ* deletion from FAPs on muscle health and function. We assessed overall muscle health by evaluating the average size of myofibers, according to our previously published pipeline (Waisman et al., 2021). Briefly, myofibers were visualized through a Phalloidin staining, segmented through Cellpose and then measured with our ImageJ plug-in LabelsToRoi. As expected, we found no differences in average myofiber size between groups (Fig. 1E). Lastly, we evaluated muscle function *in situ* via nerve mediated contractions of the TA. We found no differences in peak power or force production between FAP^no^ ^Pparγ^ mice and WTs (Fig. 1F), indicating that deletion of Pparγ in FAPs under homeostatic conditions has no effect on muscular performance.

Taken together, these data suggest that deletion of *Pparγ* within FAPs in the adult mouse has no short-term impact on systemic lipid balance, IMAT formation or muscle health under homeostatic conditions.

### IMAT formation is blocked in FAP^no^ ^Pparγ^ mice

To test whether FAP-specific deletion of *Pparγ* represses their ability to differentiate into adipocytes, we induced IMAT formation through a glycerol (GLY) injury, a widely accepted adipogenic model (Arsic et al., 2004; Joe et al., 2010; Johnson et al., 2022; Kawai et al., 1990; Kopinke et al., 2017; Mahdy et al., 2015; Mahdy et al., 2016; Norris et al., 2023; Norris et al., 2024; Uezumi et al., 2010; Waisman et al., 2021). First, we wanted to confirm successful deletion of *Pparγ* and subsequent disruption of the adipogenic program. For this, we assessed gene expression via RT-qPCR of the adipogenic signaling cascade, which is initiated through the transcription factors CCAAT/enhancer-binding proteins (C/EBPs), specifically C/EBPβ and C/EBPδ (Cristancho and Lazar, 2011; P Cornelius et al., 1994). This activates PPARγ, which in turn determines commitment to differentiation, and subsequently acts on C/EBPα (Fig. 2B). Following tamoxifen administration, TAs of WT and FAP^no^ ^Pparγ^ littermates were injured through a GLY injection and expression of these 4 adipogenic genes were assessed 3 and 5 days later (Fig. 2A). We found no difference in *C/ebpβ* or *C/ebpδ* expression between genotypes (Fig. 2B), indicating that deletion of *Pparγ* does not cause any upstream effects on the adipogenic signaling cascade. As expected, we did find a significant decrease in *Pparγ* expression in FAP^no^ ^Pparγ^ mice compared to controls (Fig. 2B), indicating successful downregulation of *Pparγ*. Additionally, we found a subsequent decrease in expression of *C/epbα,* a direct transcriptional target of PPARγ (Fig. 2B). Thus, our mouse model successfully removes *Pparγ* from FAPs and prevents the adipogenic differentiation signaling cascade from being initiated.

**Figure 2.**
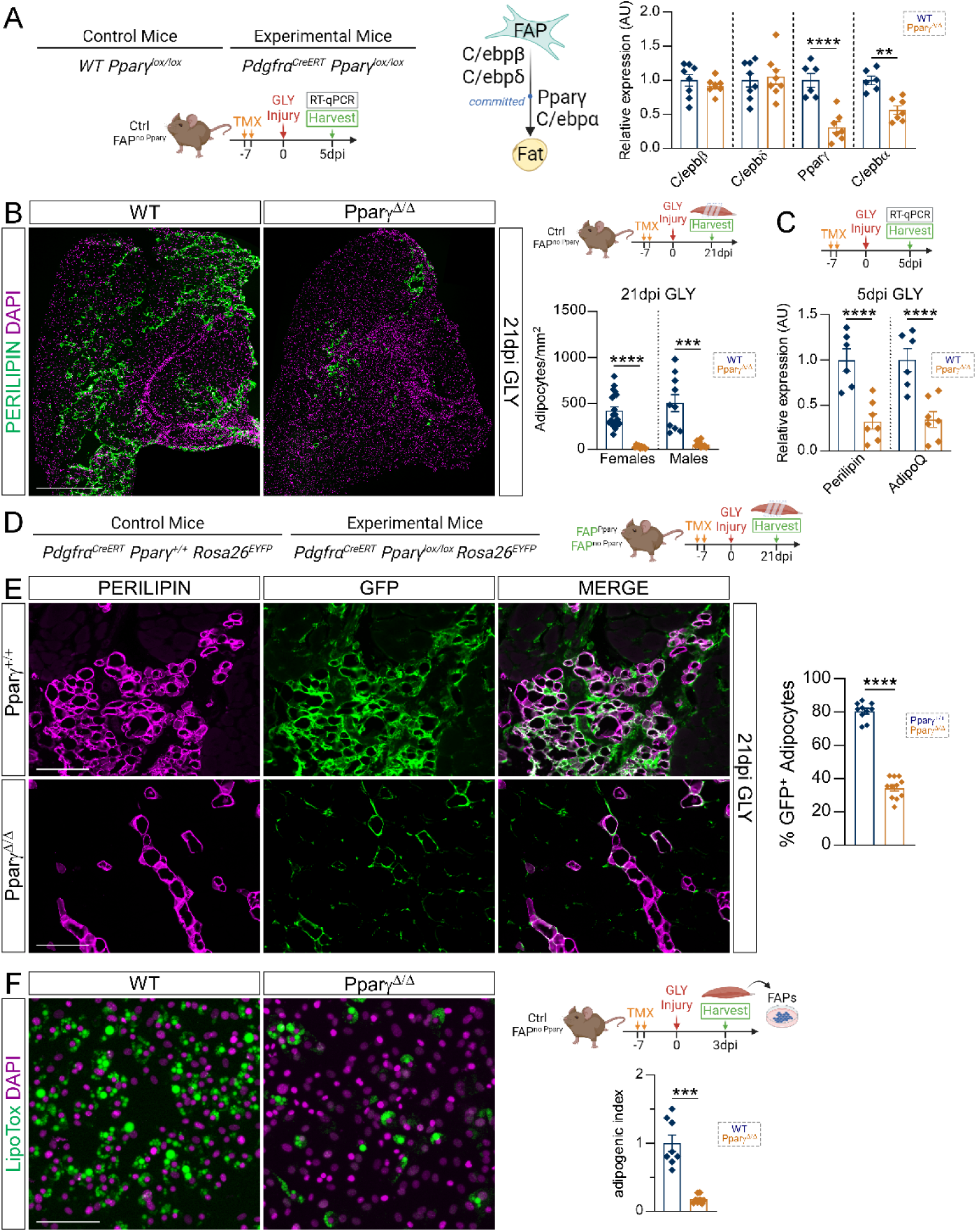
FAP^no^ ^Pparγ^ mice display limited IMAT formation following a GLY injury. **A**) (*left*) Experimental design. (*right*) RT-qPCR of early adipogenic markers in whole muscle lysate of WT (n=6-8 TAs) _and FAPno Pparγ_ (n=7-8 TAs) female mice 5 days after a GLY injury. **B)** Immuno-fluorescence of cross-section of tibialis anterior (TA) of WT and FAP^no Pparγ^ mice 21 days after a GLY injury stained for adipocytes (PERILIPIN, green). Nuclei were visualized with DAPI (magenta). Scale bar: 250µm. Quantification of number of adipocytes normalized to injured area 21 days after a GLY injury (females: WT n=20 TAs and _FAPno Pparγ n=9_ TAs; males: WT n=10 Tas and _FAPno Pparγ n=10_ TAs). **C)** RT-qPCR of late adipogenesis in whole muscle lysate 5 days after a GLY injury in female WT (n= 6 TAs) and FAP^no Pparγ^ (n=7 TAs) mice. **D)** Experimental design. **E)** Immunofluorescence of TA cross-sections of Pparγ^+/+^ and FAP^no Pparγ^ female mice 21 days after a GLY injury, stained for adipocytes (PERILIPIN, magenta) and the lineage marker GFP (green). Scale bar: 100µm. Quantification of percent of GFP^+^ adipocytes in Pparγ^+/+^ (n=10 TAs) and FAP^no Pparγ^ (n=11 TAs) mice. **F)** Immunofluorescence of *in vitro* differentiation of primary FAPs isolated from WT and FAP^no Pparγ^ mice, where lipids are visualized through a Lipidtox staining (green) and nuclei through DAPI (magenta). Scale bar: 100µm. Quantification of adipogenic differentiation between FAPs from WT (n=8 wells; n=2 mice) and FAP^no Pparγ^ (n=8 wells; n=2 mice) mice (data normalized to WT and set to 1). All data are represented as mean ± SEM. A multiple unpaired two-tailed t test followed by a Holm-Šídák post hoc test was used.

We next tested whether preventing the adipogenic signaling cascade leads to a decrease in FAP differentiation, therefore blocking IMAT formation. After tamoxifen administration and a GLY injury, TAs were harvested 21 days post injury (dpi) and IMAT formation was assessed through quantification of individual PERILIPIN^+^ adipocytes (Fig. 2C), normalized to total injured area of the TA. Excitingly, we found a strong block in IMAT formation in both female and male FAP^no^ ^Pparγ^ mice compared to littermate WTs (Fig. 2C), validating our FATBLOCK mouse model.

In addition to Cre negative controls, we initially also included Cre positive but Pparγ heterozygous animals (Pdgfrα^CreERT/+^ Pparγ^lox/+^). In these mice, tamoxifen administration should result in removal of only one allele (Pparγ^Δ/+^). When we analyzed WT, Pparγ^Δ/+^ and Pparγ^Δ/Δ^ littermates for their ability to form IMAT, we discovered that Pparγ^Δ/+^ mice displayed IMAT repression but as an intermediate phenotype between WT and FAP^no^ ^Pparγ^ mice (Supplemental Fig. 1 A & B). Combined with the fact that *Pparγ* haploinsufficiency has been previously shown to impact adipogenic differentiation *in vitro* (Rosen et al., 1999a) and in human patients (Francis et al., 2006), we only used WT mice (Pdgfrα^+/+^ Pparγ^lox/lox^) for subsequent experiments.

Next, we evaluated whether IMAT formation was prevented during early regeneration by assessing the expression of the mature adipogenic markers *Perilipin* and *Adiponectin* in whole muscle lysate 5 days after a GLY injury. There was a significant decrease in both genes in FAP^no^ ^Pparγ^ mice compared to littermate WTs (Fig. 2D) highlighting the early adipogenic repressive effect the loss of *Pparγ* has within FAPs.

To elucidate whether deletion of *Pparγ* within FAPs represses their differentiation through a cell autonomous or non-autonomous effect, we genetically labeled FAPs before injury through an EYFP reporter (Srinivas et al., 2001) and traced their differentiation into adipocytes after a GLY injury. We utilized control (Pdgfrα^CreERT/+^ Pparγ^+/+^ Rosa^EYFP^) and FAP^no^ ^Pparγ^ mice (Pdgfrα^CreERT/+^ Pparγ^lox/lox^ Rosa^EYFP^), where upon tamoxifen administration, FAPs indelibly express EYFP (Fig. 2E) (Kopinke et al., 2017). After tamoxifen administration, mice were injured with a GLY injury and harvested 21 days later. As fixation destroys endogenous EYFP expression, we counterstained EYFP with an anti-GFP antibody. We then quantified the percentage of GFP^+^ PERILIPIN^+^ adipocytes in both groups (Fig. 2F) and found a strong decrease in the percent of GFP^+^ adipocytes in FAP^no^ ^Pparγ^ mice compared to controls (Fig. 2G). Thus, deletion of *Pparγ* within FAPs causes a cell-autonomous inhibition of their differentiation into adipocytes.

To further exclude any influence from other muscle cell types on FAP differentiation, we performed an *in vitro* adipogenesis assay in isolated primary FAPs with and without *Pparγ*. We injured FAP^no^ ^Pparγ^ and WT littermates with a GLY model after tamoxifen administration, harvested TAs 3 days after injury and isolated FAPs through differential plating (Fig. 2E). Once in culture, FAPs were induced to differentiate, and adipocyte formation was measured by quantification of lipid droplets through a Lipidtox staining (Fig. 2E). We found that FAPs isolated from FAP^no^ ^Pparγ^ mice had minimal differentiation compared to WT FAPs (Fig. 2E). This further demonstrates cell autonomous repression of adipogenic differentiation without any influence of other cell types.

In addition to differentiating into adipocytes, FAPs are also progenitors of myofibroblasts, the cellular origin of fibrotic scar tissue (Joe et al., 2010; Uezumi et al., 2011). To evaluate whether prevention of FAP differentiation into adipocytes causes a shift towards a fibrotic fate, we evaluated collagen deposition in FAP^no^ ^Pparγ^ mice and WT littermates 21 days after a GLY injury through the histological stain Sirius Red (Murphy et al., 2011; Smith and Barton, 2014) (Supplemental Fig. 2D). We found no significant difference in fibrosis between genotypes in either sex (Supplemental Fig. 2D), indicating fibrosis is not affected by deletion of *Pparγ* within FAPs. Additionally, we evaluated the early fibrotic response through expression of known fibrotic genes: the known fibrotic inducer, transforming growth factor-β (*Tgfβ*) and the fibroblast marker smooth muscle α actin (*Acta2*). We found no difference in *Tgfβ* or *Acta2* expression between FAP^no^ ^Pparγ^ mice and WT littermates (Supplemental Fig. 1E). Therefore, deletion of *Pparγ* within FAPs does not redirect their fate towards a fibrotic lineage nor impact fibrotic deposition.

Taken together, deletion of *Pparγ* from FAPs causes a block in their adipogenic differentiation in a cell-autonomous manner, leading to strong repression in IMAT formation, without causing a shift in FAP fate towards a fibrotic lineage.

### Blocking FAP adipogenic differentiation changes their cellular dynamics

Due to the block in adipogenic differentiation upon removal of *Pparγ* from FAPs, we investigated how this manipulation in fate affects overall FAP dynamics during early regeneration. After tamoxifen administration, TAs of FAP^no^ ^Pparγ^ and WT mice were injured with a GLY injury and harvested throughout the early regenerative phase (Fig. 3A). We evaluated the effects in overall FAP population after deletion of *Pparγ* through quantification of the total number of FAPs (PDGFRα^+^ cells per field) at 3-, 5-, 7- and 21 days post injury (Fig. 3B). While we found no difference in total number of FAPs between genotypes 3 days after injury, there was a significant increase in the number of FAPs in FAP^no^ ^Pparγ^ mice compared to WT littermates starting at 5 dpi until 21dpi (Fig. 3B). We have previously shown that FAPs start to differentiate into adipocytes starting at 3dpi (Kopinke et al., 2017). Thus, our results indicate that preventing differentiation of FAPs into adipocytes results in a larger FAP population pool. It also provides a plausible explanation to the long-standing question on how many FAPs turn into IMAT, which, based on our result, would be ∼20%.

**Figure 3.**
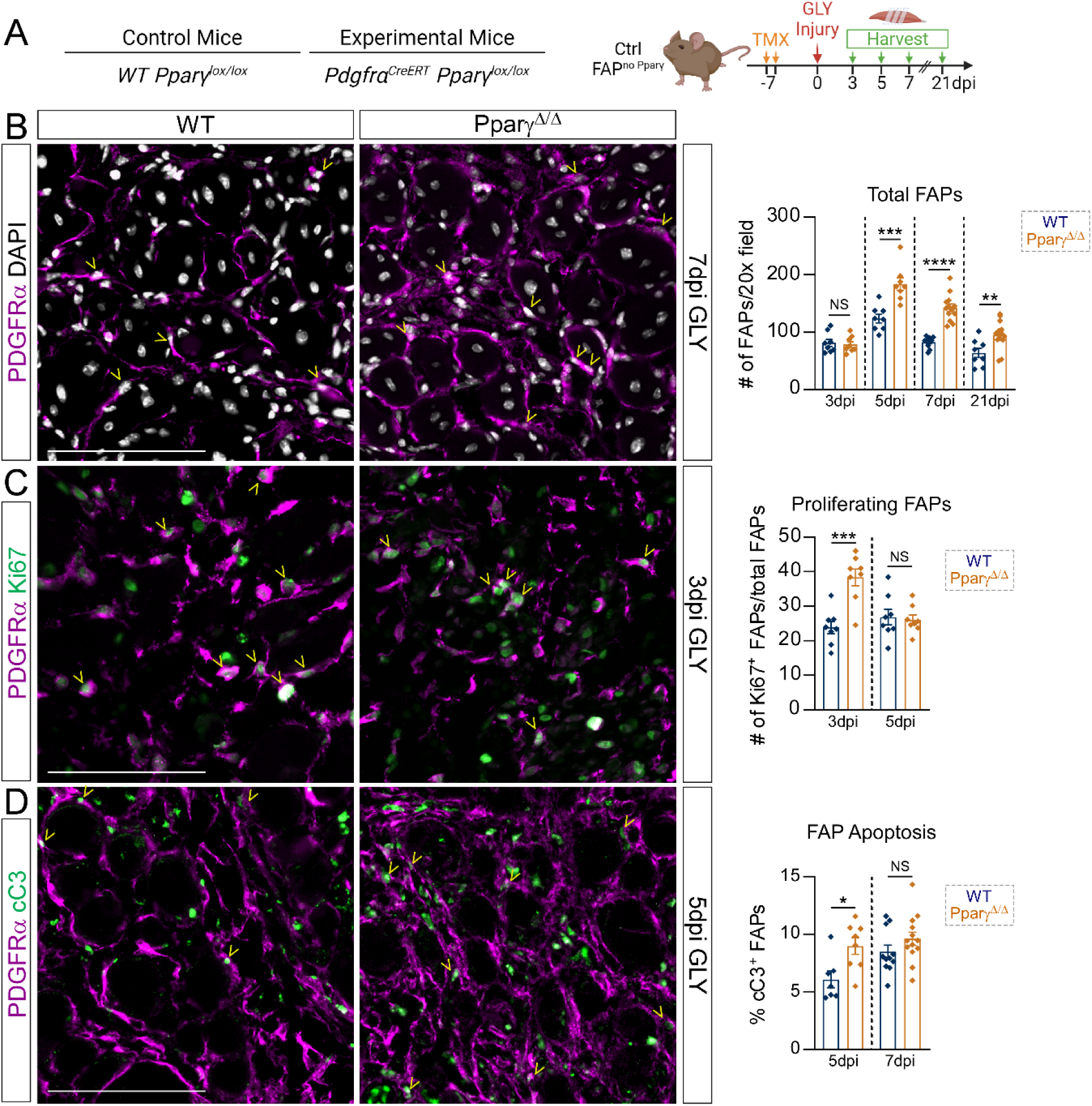
Prevention of FAP differentiation causes an increase in FAP population. **A)** Experimental design. **B)** Immunofluorescence of cross-sections of tibialis anterior (TA) muscles from WT and FAP^no Pparγ^ female mice 7 days post GLY injury stained for fibro-adipogenic progenitors (FAPs; PDGFRα, magenta) and nuclei (visualized through DAPI; white). Yellow arrowheads indicate FAPs. Scale bar: 100µm. Quantification of total FAPs per 20x field in WT (n=8-14 TAs) and FAP^no Pparγ^ (n= 8-14 TAs) female mice at 3-, 5-, 7- and 21-days post GLY injury. **C)** Immunofluorescence of TAs of WT and FAP^no Pparγ^ female mice 3 days post GLY injury, stained for FAPs (PDGFRa, magenta) and the proliferation marker, Ki67 (green). Yellow arrowheads indicate proliferating FAPs. Scale bar: 100µm. Quantification of percent of proliferating FAPs at 3- and 5-days post injury of WT (n=8 TAs) and FAP^no Pparγ^ (n=8 TAs). **D)** Immunofluorescence of cross-sections of TAs of WT and FAP^no Pparγ^ female mice 5 days post GLY injury, stained for FAPs (PDGFRa, magenta) and the canonical apoptosis marker, cleaved caspase 3 (cC3, green) 5 days post injury. Yellow arrowheads indicate FAPs that are undergoing apoptosis. Scale bar: 100µm. Quantification of percent of FAPs undergoing apoptosis in WT (n= 7-11 TAs) and FAP^no Pparγ^ (n= 8-13 TAs) mice 5- and 7-days post GLY injury. All data are represented as mean ± SEM. A multiple unpaired two-tailed t test followed by a Holm-Šídák post hoc test was used.

Due to the increase in FAP numbers, we evaluated whether deletion of *Pparγ* impacts FAP proliferation. We quantified the percentage of proliferating FAPs through the marker Antigen Kiel 67 (Ki67) within FAPs (Ki67^+^ PDGFRα^+^ cells) in FAP^no^ ^Pparγ^ and WT littermates (Fig. 3C). While we found a significant increase in proliferation rate in FAP^no^ ^Pparγ^ mice compared to WTs at 3 days post injury, both genotypes were comparable between each other 5 days after injury (Fig. 3B). This indicates that deletion of *Pparγ* within FAPs causes an acute increase in proliferation that is not sustained throughout the early regenerative phase. This could be due to the fact that in the absence of *Pparγ*, FAPs continue to proliferate, whereas FAPs with intact *Pparγ* initiate their differentiation program and cease proliferation.

Additionally, FAP apoptosis is a tightly regulated process to control FAP numbers and is significantly increased during the later regenerative phases allowing FAPs to return to pre-injury levels (Fiore et al., 2016; Lemos et al., 2015). Therefore, we quantified the percentage of FAPs (PDGFRα^+^ cells) that are undergoing apoptosis through the canonical apoptotic marker, cleaved Caspase 3 (PDGFRα^+^ cC3^+^ cells). We found an increase in percent of FAPs undergoing apoptosis in FAP^no^ ^Pparγ^ mice compared to WTs 5 days post injury (Fig. 3D). However, there was no difference between genotypes 7days post injury (Fig. 3D), depicting a transient increase in apoptosis before restoration to control levels. This transient increase in apoptosis could serve as a compensation mechanism to regulate the increased FAP population caused by prevention of their adipogenic differentiation.

Taken together, deletion of *Pparγ* in FAPs causes an increase in their population that is sustained throughout regeneration, despite an initial and transient increase in proliferation and apoptosis. Therefore, blocking adipogenic differentiation in FAPs leads to an increase in their population due to an inability to differentiate.

### IMAT impacts myofiber density and muscle function

There is a strong inverse correlation between IMAT formation and muscle mass and strength in human patients. While we and others have shown that this inverse relationship is also present in mice (Addison et al., 2014a; Addison et al., 2014b; Hilton et al., 2008; Manini et al., 2007; Norris et al., 2024), it is still unclear what the direct effects of IMAT are on muscle regeneration. Therefore, we sought to utilize our FATBLOCK mouse model to understand the direct effects of IMAT on muscle regeneration and function. Following tamoxifen administration to FAP^no^ ^Pparγ^ and WT littermates, TAs were injured with a GLY injury and after 21 days we histologically and functionally evaluated muscle regeneration (Fig. 4A).

**Figure 4.**
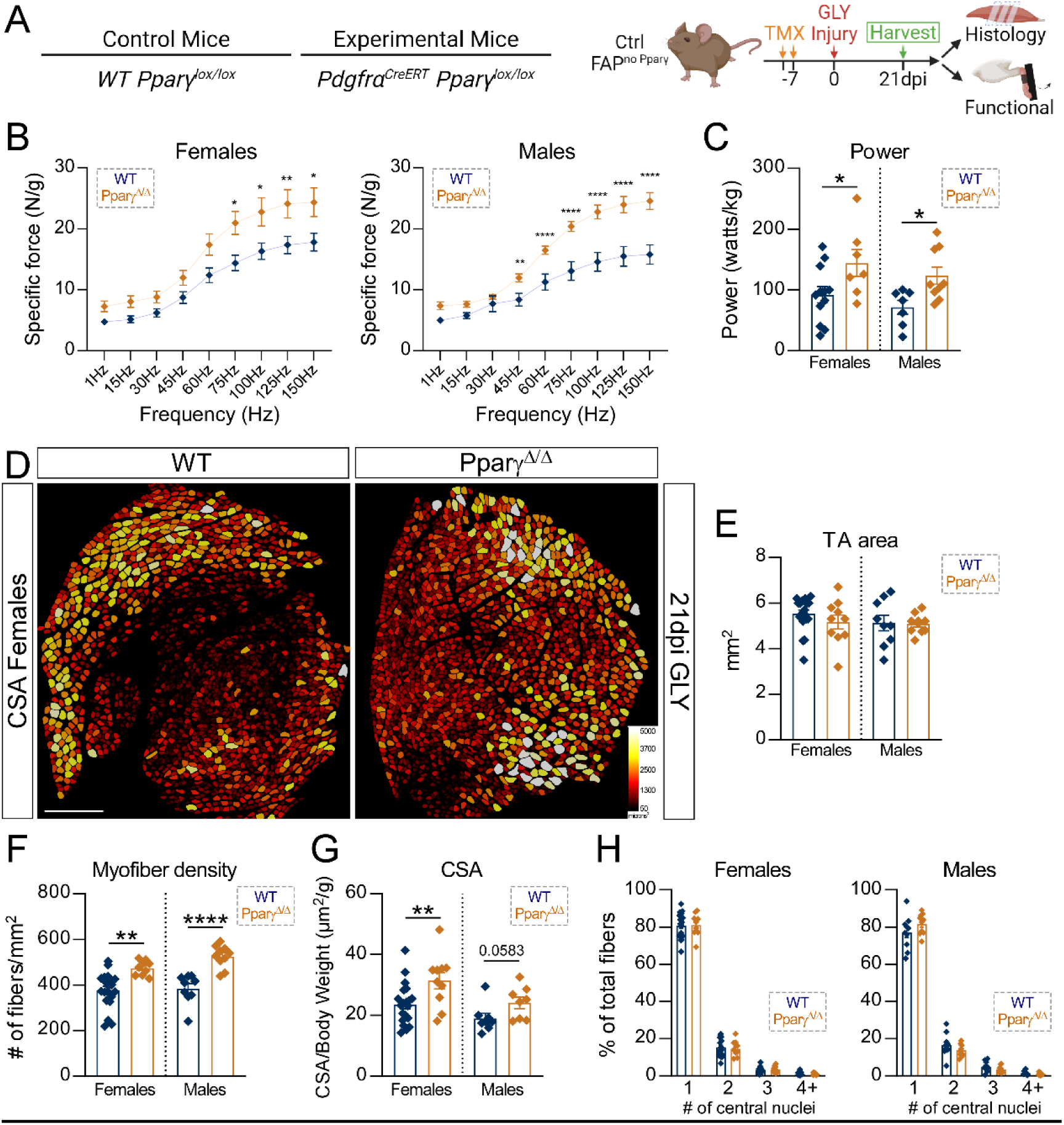
IMAT negatively impacts muscle function. **A)** Experimental outline. **(B-C)** Quantification of *in vivo* force production of the tibialis anterior (TA) muscle 21 days after a GLY injury for: **B)** Specific force generated over a range of frequencies (1-150Hz) in females (WT n=17 TAs and FAP^no Pparγ^ n=7 TAs) and males (WT n=8 TAs and FAP^no Pparγ^ n=12 TAs). **C)** Power production in females (WT n=12 TAs; FAP^no Pparγ^ n=7 TAs) and males (WT n=7 TAs; FAP^no Pparγ^ n=9 TAs). **D)** Myofibers were stained for PHAILLOIDIN followed by false color-coding according to size in WT and FAP^no Pparγ^ mice 21 days after a GLY injury. Scale bar: 500µm. **E**) Quantification of average TA cross-sectional area (CSA) 21 days after a GLY injury in females (WT n=20 TAs; FAP^no Pparγ^ n= 10 TAs) and males (WT n=9 TAs; FAP^no Pparγ^ n=10 TAs). **F)** Quantification of the number of fibers normalized to total TA cross-sectional area 21dpi GLY in females (WT n=19 TAs; FAP^no Pparγ^ n=9 TAs) and males (WT n=9 TAs; FAP^no Pparγ^ n=10 TAs). **G)** Quantification of average cross-sectional area of muscle fibers normalized to body weight in females (WT n=21 TAs; FAP^no Pparγ^ n=10 TAs) and males (WT n=8 TAs; FAP^no Pparγ^ n=8 TAs). **H)** Quantification of the number and frequency of centrally located myonuclei in females (WT n=22 TAs; FAP^no Pparγ^ n=10 TAs) and males (WT n=10 TAs; FAP^no Pparγ^ n=10 TAs). All data are represented as mean ± SEM. An unpaired two-tailed t test or a one-way ANOVA followed by a Dunnet’s or two-way ANOVA followed by a Tukey’s multiple comparison was used.

We first investigated the effects of IMAT repression on whole-muscle function in FAP^no^ ^Pparγ^ and littermate WTs (Fig. 4A). After tamoxifen and 21 days after a GLY injury to the TA, mice were anesthetized, and *in situ* force measurements were conducted on the TA muscle. Excitingly, we found a significant increase in submaximal and maximal isometric specific force and peak power output at 35% of maximal force in both sexes of FAP^no^ ^Pparγ^ compared to their WT littermates (Fig. B & C). These findings unequivocally demonstrate that IMAT negatively impacts muscle function.

To determine if the functional inhibition of IMAT is due to changes in muscle regeneration, we assessed muscle histology 21 days post GLY injury. Muscle fibers were visualized through a phalloidin staining, segmented and measured through our previously published pipeline (Fig. 4D). We found no difference in total TA area between genotypes (Fig 4E), indicating that prevention of IMAT formation is not causing any gross phenotypic differences. However, we discovered a significant increase in the density of myofibers (number of myofibers normalized to TA area) in FAP^no^ ^Pparγ^ mice compared to WT littermates in both sexes (Fig. 4F), indicating that IMAT is negatively affecting the number of fibers that are able to regenerate after an injury. Next, we assessed the average cross-sectional area (CSA) of myofibers and found that FAP^no^ ^Pparγ^ mice had a higher average CSA compared to WTs (Fig. 4G). As we have seen an increase in myofiber density in both sexes and an increase in average CSA in females, we sought to explore whether myoblast fusion could be affected by IMAT. We assessed myoblast fusion by quantifying the percentage of centrally located nuclei in regenerated myofibers. We found no difference in the distribution of the percentage of number of centrally located nuclei (Fig. 4H), indicating that IMAT has no effect on myoblast fusion.

Additionally, we evaluated IMAT and muscle regeneration following a cardiotoxin (CTX) injury model, where IMAT formation is limited in comparison to a GLY injury (Kopinke et al., 2017; Norris et al., 2023; Norris et al., 2024). Following tamoxifen administration to FAP^no^ ^Pparγ^ and WT littermates, TAs were injured with CTX and harvested 21 days after injury (Supplemental Fig. 2A). We found a significant decrease in IMAT formation in FAP^no^ ^Pparγ^ females compared to WT littermates although the total IMAT amount was significantly less (∼20-fold) compared to a GLY injury (Supplemental Fig. 2B). However, we found no difference in IMAT between genotypes in males (Supplemental Fig. 2B). Whether this indicates sex-specific differences in IMAT formation after a CTX injury or we approached a threshold for how far we can repress IMAT formation remains to be seen. We next evaluated muscle regeneration, as previously mentioned, by assessing myofiber density and average CSA (Supplemental Fig. 2C-E). We found no difference in any of these measurements between genotypes in either sex, indicating that deletion of *Pparγ* within FAPs has no effect on muscle regeneration after a CTX injury.

Taken together, our results demonstrate that IMAT, if allowed to form, causes a decline in myofiber density and size resulting in a functionally impaired muscle. In addition, the fact that IMAT only had an impact on myofiber regeneration post GLY injury, which causes dramatically more accumulation of adipocytes than CTX, indicates the presence of an IMAT threshold that is needed for IMAT to have a negative impact.

### IMAT physically blocks the formation of new fibers

To determine how IMAT might affect myofiber density and size, we evaluated the early myogenic process in FAP^no^ ^Pparγ^ and control littermates. Following tamoxifen administration, TAs were injured with a GLY injury and were harvested at 3-, 4-, 5- and 7 days post injury (Fig. 5A). We first analyzed the expression values of the MuSCs marker *Pax7* and another myoblast marker, *MyoD*, by RT-qPCR in FAP^no^ ^Pparγ^ and WT littermates (Fig. 5B). We found no difference in expression of *Pax7* or *MyoD* between genotypes 3- and 5 days post GLY injury (Fig. 5B), indicating that deletion of IMAT does not affect early myogenic differentiation. Next, we quantified the total number of myoblasts, characterized by Myogenin expression (MyoG^+^ cells).

**Figure 5.**
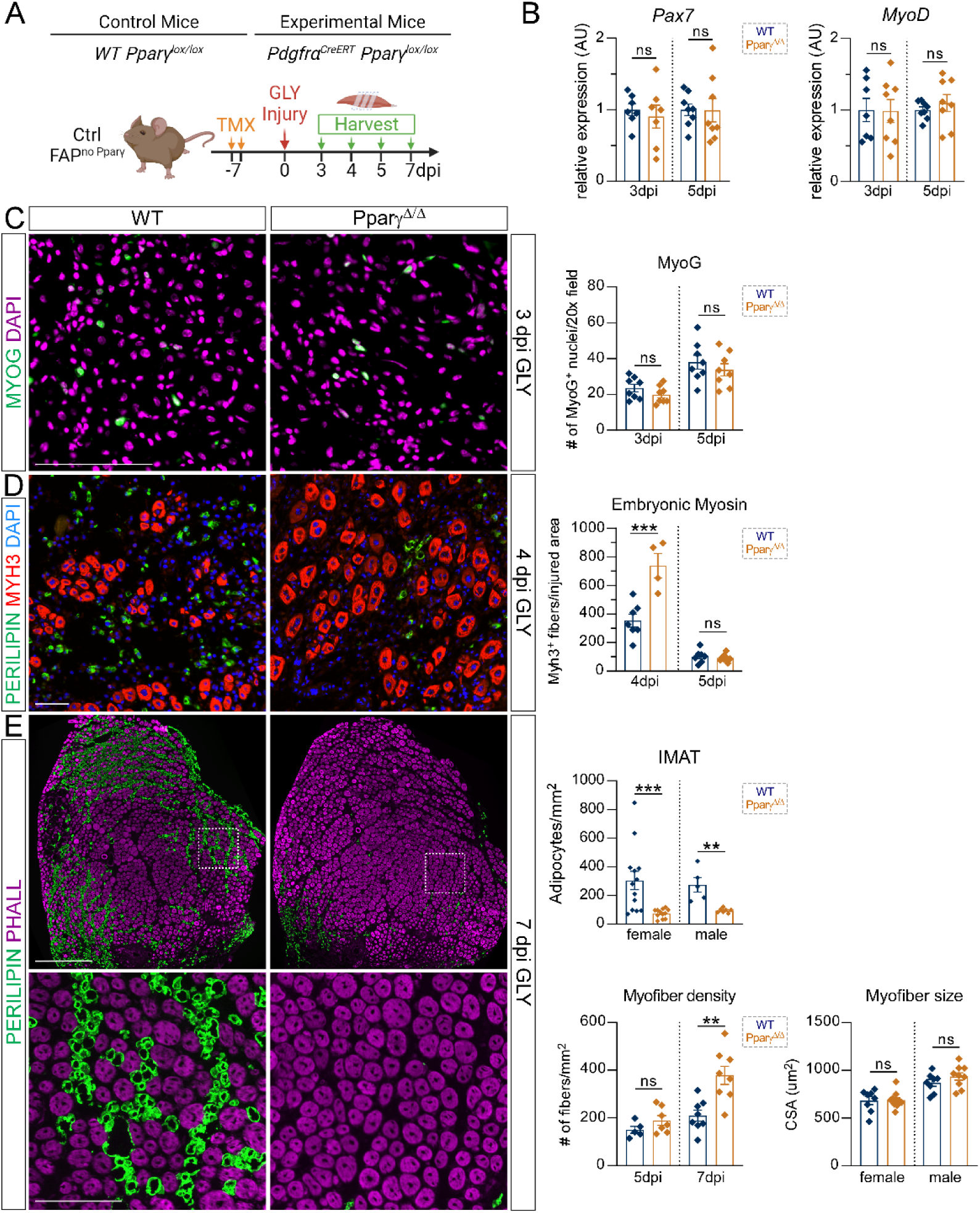
IMAT limits myofiber density but not myogenic differentiation. **A)** Experimental outline. **B)** RT-qPCR of whole tibialis anterior (TA) muscle lysate from WT (n=6-7 TAs) and FAP^no Pparγ^ (n=7-8 TAs) female mice 3- and 5 days post a GLY injury for *Pax7* (MuSCs) and *MyoD* (myoblasts). **C)** Immuno-fluorescence of TA cross-sections from WT and FAP^no Pparγ^ mice 3 days post GLY injury, stained for MYOG (green; myoblasts) and nuclei visualized through DAPI (magenta). Scale bar: 100µm. Quantifications of MYOG^+^ nuclei per 20x field of view in WT (n=7-8 TAs) _and FAPno Pparγ_ (n=8 TAs) 3- and 5 days post GLY injury. **D)** Immuno-fluorescence detection of newly formed myofibers (marked by embryonic myosin; MYH3, red), adipocytes (PERILIPIN, green) 4 days post GLY injury in WT and FAP^no Pparγ^ mice. Nuclei were visualized through DAPI (blue). Scale bar: 100µm. Quantification of MYH3^+^ myofibers normalized to injured area in WT (n=7 TAs) and FAP^no Pparγ^ (n=4-8 TAs) female mice 4- and 5 days post GLY injury. **E)** Immunofluorescence of TA cross-section from female WT and FAP^no Pparγ^ mice 7 days post GLY injury stained for myofibers (PHALLOIDIN, magenta) and adipocytes (green). Scale bar: 500µm. Zoom Scale bar: 150µm. Quantification of adipocytes normalized to injured area 7 dpi GLY in females (WT n=13 TAs; FAP^no Pparγ^ n=12 TAs) and males (WT n=5 TAs; FAP^no Pparγ^ n=7 TAs). Quantifications of number of myofibers normalized to injured area in female WT (n=5-8 TAs) and FAP^no Pparγ^ (n=7-8 TAs) mice 5- and 7-days post GLY injury. Quantification of average cross-sectional area (CSA) of myofibers in females (WT n=8 TAs; FAP^no Pparγ^ n=11 TAs) and males (WT n=8 TAs; FAP^no Pparγ^ n=9 TAs). All data are represented as mean ± SEM. An unpaired two-tailed t test was used.

At 3- and 5 days post GLY injury, we found no difference in MyoG^+^ myoblasts between FAP^no^ ^Pparγ^ and WT littermates at either timepoints (Fig. 5B). Thus, IMAT does not impact myogenic differentiation.

We next evaluated newly formed fibers, characterized by expression of embryonic myosin (MYH3^+^ fibers), in FAP^no^ ^Pparγ^ and WT littermates at 4- and 5-days post GLY (Fig. 5D). We detected a robust increase in the number of MYH3^+^ fibers 4 days post GLY in FAP^no^ ^Pparγ^ compared to WT littermates (Fig. 5D). By 5dpi, however, only few MYH3^+^ fibers can still be detected with no difference between genotypes (Fig. 5D), indicating a dramatic but transient increase in newly formed fibers. We also stained for adipocytes and can already detect PERILIPIN^+^ adipocytes at day 4 in control mice. In contrast, only few adipocytes formed in the FAP^no^ ^Pparγ^ mice. Combined, this indicates the exciting possibility that IMAT might act as a physical barrier, thereby repressing fiber formation.

Next, we looked at day 7 post GLY, a time point where most myofibers have regenerated but still need to undergo hypertrophy to reach pre-injury size (Fig. 5E). Similar to 21dpi (Fig. 2), we see robust IMAT infiltration, which is blocked upon loss of *Pparγ* (Fig. 5F). When we quantified the number of myofibers present per injured area, we detected no difference between genotypes at 5 days post GLY injury but found a dramatic increase in myofiber density 7 days post injury in FAP^no^ ^Pparγ^ mice compared to control littermates (Fig. 5G). In contrast, myofiber size, as assessed by CSA, remains unchanged at both timepoints.

Taken together, our results demonstrate that IMAT does not affect myogenic differentiation but rather acts as a physical barrier that blocks myofiber formation. Additionally, these data also strongly argue that IMAT also physically prevents post-regenerative hypertrophy needed for myofibers to return to their pre-injury size by limiting the space the myofibers can grow into.

### IMAT does not act as a local signaling center

IMAT has also been shown to secrete anti-myogenic signals (Coles, 2016; Collins et al., 2022; Li et al., 2017; Sachs et al., 2019). Therefore, we evaluated a variety of signaling pathways that are known to influence muscle regeneration. One crucial adipogenic signal that influences myofiber size is Leptin (Collins et al., 2022). To test whether IMAT affects systemic and local Leptin concentrations, we measured circulating Leptin levels in serum and the TA 5 days post GLY injury through an ELISA (Fig. 6A). We found no differences in Leptin concentration in circulating serum or TA lysate between FAP^no^ ^Pparγ^ and control mice.

**Figure 6.**
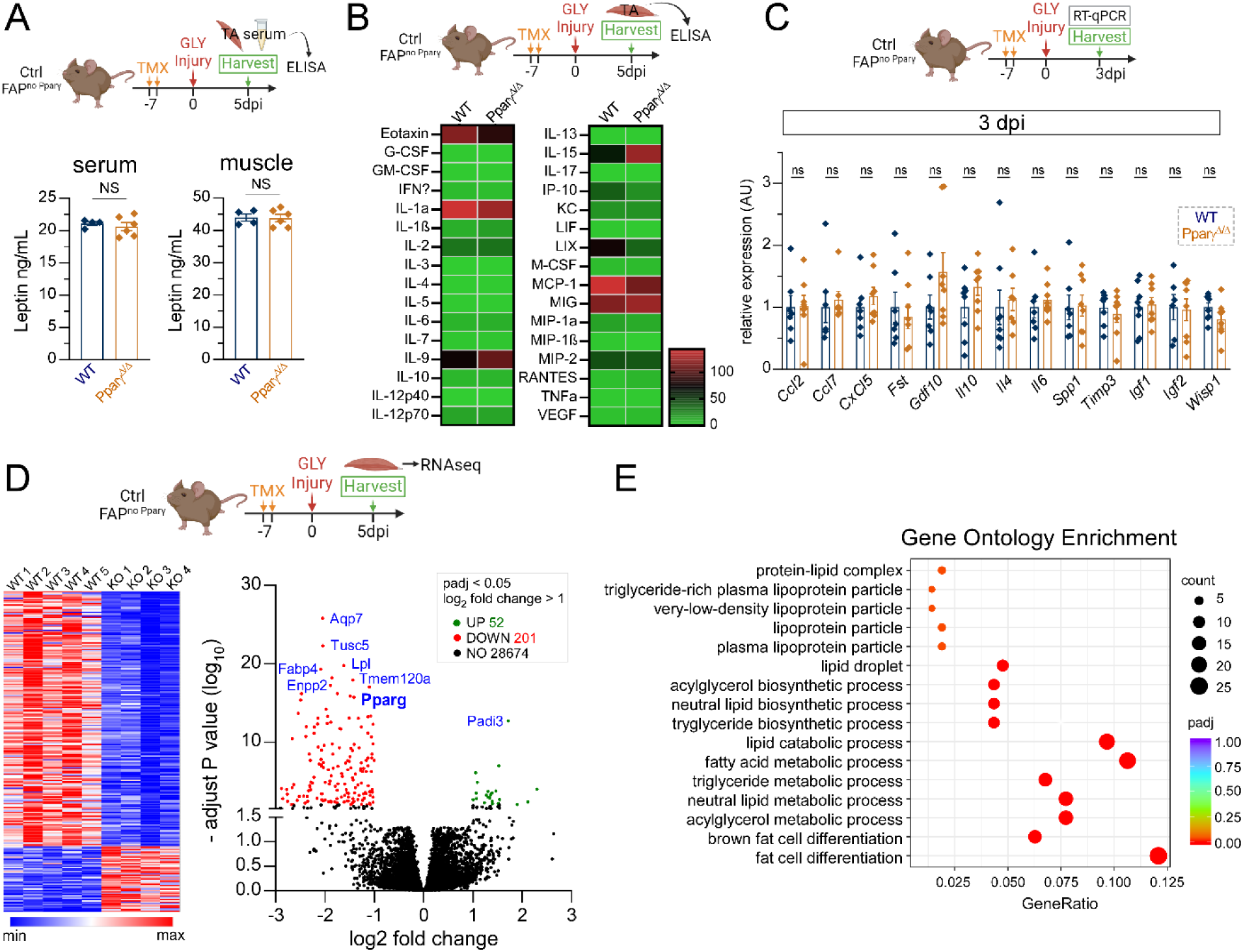
IMAT does not cause aberrant signaling during regeneration. **A)** Experimental outline. Quantifications of circulating leptin concentration in serum and tibialis anterior (TA) lysate from WT (n=4 mice/TAs, respectfully) and FAP^no Pparγ^ (n=6 mice/TAs) female mice 5 days post GLY injury. **B)** Experimental outline. Multiplex enzyme-linked immunosorbent assay (ELISA) of multiple cytokines from whole protein lysate of TAs of female WT (n=5 TAs) and FAP^no Pparγ^ (n=5 TAs) mice 5 days post GLY injury. **C)** Experimental outline. RT-qPCR of whole muscle TA lysate of female WT (n= 6-8 TAs) and FAP^no Pparγ^ (n= 6-8 TAs) mice 3 days post GLY injury for FAP-derived signals. **D)** (*left)* Heatmap of transcriptomics data generated through bulk RNA sequencing (RNAseq) of whole muscle TA lysates from female WT (n= 5 TAs) and FAP^no Pparγ^ (n=4 TAs) mice 5 days post GLY injury. (*right*) Volcano plot of differentially expressed genes with Log2 fold change ≥ 1 and an adjusted p-value of < 0.05 (genes in red are down- and in green up-regulated). **E)** Gene ontology enrichment results for differentially expressed genes between WT and FAP^no Pparγ^ mice. All data are represented as mean ± SEM. An unpaired two-tailed t test or a multiple unpaired two-tailed t test followed by a Holm-Šídák post hoc test was used.

We then asked if loss of IMAT has any effects on cytokine secretion (Fig. 6B). For this, we ran muscle-specific protein lysates through a 32-Plex Cytokine assay. Similar to Leptin, IMAT repression had no significant impact.

We and others have shown that FAPs are crucial producers of many pro-myogenic cues (Contreras et al., 2021; Flores-Opazo et al., 2024; Kopinke et al., 2017; Norris et al., 2023). To test whether loss of *Pparγ* itself or the increase in FAPs in the FAP^no^ ^Pparγ^ mice might cause any differences, we screened multiple myogenic cues by RT-qPCR at 3 (Fig. 6C) and 5 days post GLY injury (Supplemental Fig. 3A). Again, we detected no differences between genotypes highlighting that deletion of *Pparγ* does not result in a discernable secretory phenotype.

Last, we aimed to objectively discover any potential differences by comparing the transcriptional profile of FAP^no^ ^Pparγ^ to control mice. For this, we isolated whole muscle RNA from TAs 5 days post GLY injury and performed bulk RNA sequencing (Fig. 6D). To our surprise, we only detected ∼200 down and ∼50 up regulated genes (Log2 fold change >1 and padj <0.05). Besides *Pparγ*, most of the top down-regulated genes, such as *Fabp4, Lpl, Tusc5, Tmem120a, Aqp7* and *Enpp2*, have known roles in adipogenesis (Beaton et al., 2015; Frühbeck et al., 2023; Gonzales and Orlando, 2007; Prentice et al., 2019; Qian et al., 2022). Fittingly, performing a gene ontology enrichment analysis highlighted that most processes, which changed upon *Pparγ* deletion, are related to adipogenesis and fat metabolism (Fig. 6E). One of the hits suggested that brown fat cell differentiation might be affected. However, we failed to confirm any differences in multiple brown fat specific genes by RT-qPCR at 7 days post GLY (Supplemental Fig. 3B).

Thus, removal of *Pparγ* from FAPs and the subsequent repression of IMAT formation does not cause any detectable changes in adipokines, cytokines or myogenic factors further strengthening our argument that IMAT solely acts as a physical barrier.

## DISCUSSION

Accumulation of IMAT is a shared feature of age and disease and is strongly associated with decreased muscle strength and function (Burakiewicz et al., 2017; Flores-Opazo et al., 2024; Goodpaster et al., 2005; Goodpaster et al., 2003; Goodpaster et al., 2006; Goodpaster et al., 2000a; Goodpaster et al., 2000b; Hogarth et al., 2019; Milad et al., 2017; Murphy et al., 1986b; Norris et al., 2024; Willcocks et al., 2016; Wokke et al., 2014). However, it is still unresolved whether IMAT is actively repressing muscle regeneration and/or function. In this study, we created an inducible IMAT prevention mouse model, called FATBLOCK, where we can inhibit the cellular origin of IMAT, FAPs, from differentiating into adipocytes. Through deletion of peroxisome proliferator-activated receptor γ (*Pparγ)*, the master regulator of adipogenic fate (Rosen et al., 2000), from FAPs, their adipogenic differentiation was blocked in a cell autonomous manner with no shift in their cellular fate. Upon an adipogenic injury caused by GLY, we found that IMAT decreases the density and size of regenerated myofibers resulting in a potent boost in the functional recovery of muscle. Interestingly, the repressive impact of IMAT was independent of post-regenerative MuSC-driven myogenesis. In addition, IMAT did not act as a major signaling center itself or broadly influenced intercellular communication. Instead, our data reveal that IMAT functions as a physical barrier: early, to prevent nascent myofibers from forming, and later, to restrict their post-regenerative hypertrophic growth. Together, our results define the negative impact of IMAT on regenerating muscle and emphasize the importance of developing novel approaches to target IMAT shortly after an adipogenic injury to prevent long-term muscle loss.

### IMAT: innocent bystander or active participant?

Several studies have tried to address whether IMAT might have a direct influence on muscle with mixed results. The Feige group took advantage of mice with a whole-body knockout of peroxisome proliferator-activated receptor γ (*Pparγ)*, the master regulator of adipogenic fate (Rosen et al., 2000). These mice are incapable of developing adipose tissue including intramuscular fat. Confirming this, *Pparγ ^-/-^*mice did not form intramuscular fat 14 days post GLY injury. Importantly, *Pparγ ^-/-^* mice displayed compromised myofiber recovery suggesting that IMAT is important for muscle regeneration (Dammone et al., 2018). However, *Pparγ ^-/-^*mice are lipodystrophic, as they are completely devoid of all fat, and develop liver steatosis resulting in severe metabolic defects. In addition, PPARG functions in other cell types, including MuSCs and macrophages (Singh et al., 2007; Varga et al., 2016). Thus, the observed phenotype could be due to severe metabolic distress and/or off-target effects.

In contrast, using a clever and rigorous cell ablation approach to kill adipocytes, the Kuang group found that fewer myofibers regenerated after a CTX injury when intramuscular fat formation was repressed (Liu et al., 2012). However, the Cre line used to ablate adipocytes is also active in endothelial cells (Cataltepe et al., 2012; Drummond et al., 2018; Elmasri et al., 2009; Hatley et al., 2012), macrophages (Hui et al., 2010; Maeda et al., 2005; Makowski et al., 2005) and adipogenic progenitors (Shan et al., 2013) and, thus, would have ablated these cell types as well. In addition, they only used a CTX injury, where we found myofiber regeneration to be independent on IMAT (Supplemental Fig. 2).

Using another elegant fat-ablation model, the Meyer group showed that intramuscular fat causes contractile tension deficits of TA muscles 21 days post GLY injury (Biltz et al., 2020). However, like the germline *Pparγ* KO model, this fat-ablation model also causes lipodystrophy. This resulted in lower cross-sectional area and peak tetanic tension at baseline, thereby providing an alternative explanation for the observed phenotype. The Meyer group recently demonstrated that the baseline deficits are due to disrupted Leptin levels, a potent adipokine secreted by fat required for the development of normal muscle mass and strength (Collins et al., 2022). We failed to detect any differences in myofiber size or muscle function prior to injury (Fig. 1), most likely due to preserved Leptin levels (Fig. 5A).

Here, we describe our new FATBLOCK model in which we genetically deleted *Pparγ* specifically and postnatally in FAPs to inhibit their adipogenic differentiation into IMAT without causing major phenotypic changes. In the absence of injury, 15-week-old FAP^no^ ^PPARG^ mice displayed normal body weights and showed no indication of lipodystrophy 5 weeks post tamoxifen administration. Importantly, postnatal deletion of *Pparγ* specifically in FAPs did not cause massive cell death nor had it any impact on muscle homeostasis, as assessed by muscle weight, function and myofiber size (Fig. 1). Thus, our model allowed us to conclusively and irrefutably test the impact of IMAT on muscle function particularly during regeneration. While we cannot rule out that IMAT might also cause contraction deficit (Biltz et al., 2020), our results demonstrate that IMAT functionally impairs muscle function by acting as a physical barrier, resulting in a muscle that is comprised of fewer myofibers that are of smaller size.

### IMAT threshold: what are the implications?

We recently compared IMAT formation to myofiber regeneration between CTX and GLY injuries and found that IMAT is strongly but negatively correlated with myofiber size in that the more IMAT a muscle had formed, the smaller the newly regenerated myofibers were (Norris et al., 2024). Fittingly, here we only detected the negative influence of IMAT on myofiber regeneration after a GLY but not a CTX injury. Given the significant difference in IMAT levels between the two injuries, our results strongly argue for the existence of an IMAT threshold, and that, only when exceeded, will IMAT become detrimental to muscle regeneration. The existence of an IMAT threshold could have major implications for murine models that try to mimic human conditions, which display massive IMAT infiltration as a key pathological phenotype. For example, in our previous study, we demonstrated that IMAT formation is dependent on the genetic background (i.e. strain) and sex of the mouse (Norris et al., 2024). Female mice across different strains and injuries produced more IMAT, while C57/BL6 mice were extremely lean and formed very little IMAT. Therefore, if a specific IMAT threshold is necessary for IMAT to influence myofiber regeneration, careful consideration must be given to the sex, strain, and injury type when studying the effects of IMAT on muscle health.

### Removal of *Pparγ* has no major impact on FAPs

FAPs are crucial during the repair of damaged muscles by secreting beneficial factors (Joe et al., 2010; Kopinke et al., 2017; Lukjanenko et al., 2019; Murphy et al., 2011; Santini et al., 2020; Uezumi et al., 2010; Uezumi et al., 2021; Uezumi et al., 2011; Wosczyna et al., 2019). Therefore, we carefully assessed any potential impact of the loss of *Pparγ* on the health of FAPs. Overall, we detected a ∼20% increase in the overall FAP population in FAP^no^ ^Pparγ^ mice starting at day 5 and persisting until our end point of 21 days post injury. While we noticed a modest increase in FAP proliferation at day 3 and a slight increase in apoptosis at day 5, we believe that the persistent increase in FAP numbers is rather due to the fact that FAPs lacking *Pparγ* fail to differentiate into adipocytes and thereby maintain their FAP fate. This also suggests that around 20% of all FAPs eventually differentiate into adipocytes following a GLY injury. It will be interesting to determine whether these 20% are a distinct pro-adipogenic subpopulation or whether every FAP has the potential to form fat but only one in five actually does so.

The observed increase in FAPs could result in an increase in pro-myogenic factors, thereby providing an alternative explanation for why loss of FAP *Pparγ* could improve muscle regeneration. After carefully assessing multiple known factors, we failed to detect any differences in FAP^no^ ^Pparγ^ mice compared to controls. In fact, we were unsuccessful in detecting any changes in a variety of signals (adipokines and cytokines) or global transcriptional changes beyond genes involved in adipogenesis. Together, this further argues for IMAT physically impacting muscle regeneration.

Besides adipocytes (Hogarth et al., 2019; Joe et al., 2010; Kopinke et al., 2017; Liu et al., 2012; Scott et al., 2019; Stumm et al., 2018; Uezumi et al., 2010; Uezumi et al., 2011), FAPs can also differentiate into myofibroblasts, the cellular origin of tissue fibrosis (Joe et al., 2010; Uezumi et al., 2011). Moreover, an increase in FAP numbers is often associated with an increase in fibrosis (Joe et al., 2010; Uezumi et al., 2011). However, despite the increase in FAPs, we fail to detect any increase in fibrotic scar tissue formation. Instead, FAPs remain as PDGFRA-expressing FAPs excluding the possibility that FAPs differentiate into myofibroblasts as an alternative fate upon *Pparγ* deletion.

### IMAT’s role in acute injuries vs chronic conditions

One important aspect to consider is how IMAT arises and whether it would have the same effect on muscle depending on etiology of the disease. In this study, we have shown that in an injury setting, which allows IMAT to form while muscle is regenerating, it will hinder the regenerative process, leading to a loss in muscle fiber density and, ultimately, muscle function. One example of this type of acute but adipogenic injury is rotator cuff tears in the supraspinatus muscle. The greater the tear, the more FAPs expand, leading to massive IMAT infiltration (Gladstone et al., 2007; Goutallier et al., 2003; Klatte-Schulz et al., 2014; Liu et al., 2016; Trudel et al., 2019). As a consequence, muscle regeneration is compromised, resulting in muscle atrophy and declined functional output. Thus, preventing this rapid IMAT infiltration could have massive benefits.

However, the effects of IMAT on muscle could be substantially different in other diseases devoid of injury, where metabolic defects lead to progressive IMAT infiltration. For example, during ageing and obesity, IMAT progressively infiltrates muscle over time and its effects on muscle could be substantially different to an acute injury (Goodpaster et al., 2005; Goodpaster et al., 2003; Goodpaster et al., 2006; Goodpaster et al., 2000a). Adipose tissue is a potent endocrine organ, and IMAT in particular has been shown to modulate muscle insulin sensitivity, possibly via secretion of inflammatory cytokines (Sachs et al., 2019). There is also some evidence that adipocytes could impair myogenesis *in vitro* via secretion of certain cytokines such as IL6, IL-1β and TNF-α (Pellegrinelli et al., 2015; Seo et al., 2019; Takegahara et al., 2014), or myofiber size *in vivo* through LEPTIN (Collins et al., 2022). In our IMAT prevention model, we found no differences in these targets, as well as other inflammatory markers, arguing that injury-induced IMAT in a metabolically healthy organism results in a significantly different response.

Neuromuscular diseases such as Duchenne muscular dystrophy (DMD) are known to have bouts of degeneration followed by regeneration of muscle (mimicking injury), while also having dysfunctional signaling and metabolism (Dabaj et al., 2021; Xu et al., 2023), seen by abnormal amino acid, energy, and lipid metabolism, as well as calcium balance, mitochondrial function and insulin resistance that is independent of corticosteroid use (Rodríguez-Cruz et al., 2015). Thus, DMD, and similar diseases, might represent a scenario where IMAT might act both as physical barrier to prevent the regeneration of damaged myofibers but also as a potent immune modulator. Unfortunately, the lack of a suitable mouse model, which displays progressive but complete replacement of muscle with IMAT, prevents us from applying our FATBLOCK model to define the long-term impact of IMAT on muscle health in chronic conditions.

In sum, this work reveals the novel finding that acute infiltration of IMAT physically limits the formation of new myofibers and restricts their post-regenerative hypertrophic growth. As a result, the regenerated muscle is functionally weakened. Our work also highlights the importance of blocking IMAT formation as a potent strategy to improve the healing of acute but adipogenic injuries such as rotator cuff tears.

## MATERIALS AND METHODS

### Animal Studies

The alleles utilized have been previously described. To genetically target fibro-adipogenic progenitors (FAPs), we utilized the *Pdgfrα^CreERT2^* allele (Chung et al., 2018). To prevent FAPs from differentiating into adipocytes, we genetically deleted the master regulator of adipogenesis, *Pparγ*, through the allele B6.129-Pparγ^tm2Rev^/J (He et al., 2003). To obtain a mouse model with high IMAT formation, we backcrossed both alleles onto a 129S1/SvlmJ strain for 3 generations. Based on our previous work, the 129S1/SvlmJ strain has a higher adipogenic potential compared to the C57BL/6J strain. We accomplished deletion of *Pparγ* from FAPs in adult 10-13 week old mice upon Tamoxifen administration (TRC, T006000) dissolved in corn oil and administered through oral gavage for two consecutive days, at a dose of 200-250 mg/kg to all mice. Muscle injuries were performed a week after the last tamoxifen treatment to allow metabolic clearance of tamoxifen from mice. Littermates were used for each timepoint and experiment; mice lacking the *Cre* allele were used as controls. Littermates with the *Cre* allele and heterozygous or homozygous for the *floxed* allele were analyzed separately. Upon observation of a haploinsufficiency of the *Pparγ* gene for IMAT formation, we utilized littermates lacking the *Cre* allele (Pdgfrα^+/+^; Pparγ^fl/+^ or Pparγ^fl/fl^) as controls, and mice with the *Cre* allele that are homozygous for the floxed allele (Pdgfrα^CreERT2/+^; Pparγ^fl/fl^) as the experimental group. Mice were housed in standard ventilated cages at a controlled temperature (22-23 °C), with 40-50% humidity and ad libitum access to food and water. All animal work was approved by the Institutional Animal Care and Use Committee (IACUC) of the University of Florida.

### Muscle injuries

All injuries were performed on adult 10-13 week old mice. Mice were put under anesthesia with isoflurane and the Tibialis Anterior (TA) was injected with 30-50 µL of 50% glycerol (GLY; Acros Organics, 56-81-5) or with 10 µM cardiotoxin (CTX; Naja Pallidum, Lotaxan, L8102-1MG), both diluted in sterile saline.

### *In situ* force production

Functional testing of the Tibialis Anterior muscle (TA) was measured *in-situ* by stimulation of the sciatic nerve as previously described (Dong G, 2023; Palzkill et al., 2024a; Palzkill et al., 2024b; Spinazzola et al., 2015). Briefly, Mice were anaesthetized with an intraperitoneal injection of xylazine (1 mg/kg) and ketamine (10 mg/kg) and subsequent doses of ketamine were given as needed for maintenance. The TA was isolated from its distal insertion and the distal tendon was secured with a 4-0 silk suture and attached to the lever arm of a force transducer (Cambridge Technology, Model No. 2250). Muscle contractions were induced by stimulating the sciatic nerve with bipolar electrodes using 0.02 ms square wave pulses (Aurora Scientific, Model 701 A stimulator). Data collection and servomotor control were managed using the Lab View-based DMC program (version 615A.v6.0, Aurora Scientific Inc). After determining the optimal muscle length through twitch contractions, a force frequency curve was performed. Isometric contractions were elicited using 500ms train (current 2mA, pulse width 0.2ms) at stimulation frequencies of 1Hz, 15Hz, 30Hz, 45Hz, 60Hz, 75Hz, 100Hz, 125Hz, and 150Hz with one minute of rest between contractions. Peak twitch and tetanic force levels were reported as absolute and specific (normalized to muscle weight) force levels. Maximum isometric force was determined from the peak tetanic force elicited from the force-frequency analysis. Given most practical uses of skeletal muscle require an aspect of muscle shortening to generate power, a single after-loaded isotonic contraction was performed against a predetermined, submaximal load. This involved electrically stimulating the nerve with supramaximal voltage (2mA, 0.2ms pulse width, 100ms train duration) at 150Hz allowing the muscle to shorten once it reaches 35% of maximum isometric tension. This allows for the quantification of muscle performance characteristics including force (newtons), displacement (mm), shortening velocity (m/s), and mechanical power (watts). Shortening velocity was calculated as the change in distance (mm) from a 10ms period which began 20ms after the initial length change. Peak power was calculated as the product of the shortening velocity (m/s) and corresponding force (N/kg).

### In vitro studies

To isolate FAPs, TAs from mice lacking the Cre allele (WT) and Pdgfrα^CreERT2/+^; Pparγ^fl/fl^ (FAP^no^ ^Pparγ^) littermates were injured with GLY (as described above). TAs were harvested 3 days after injury and FAPs were isolated through differential plating as described previously (Norris et al., 2023). Briefly, both TA muscles were mechanically and enzymatically digested in DMEM (Gibco, 10566024), 1% Pen/Strep, 50 mg/mL collagenase IV (Worthington, LS004188), and 6 U/mL Dispase (Gibco, 17105041) for 1 h at 37 °C with constant agitation, until a mostly homogenous unicellular suspension was obtained. This suspension was then filtered (70µm filter; Fisher 22363548) and plated in DMEM at 37 °C for 30min to allow FAPs to adhere to the plate. The supernatant was then removed, and the plate was gently washed with warm PBS 3 times, obtaining isolated FAPs and cultured in DMEM (Gibco, 10566024) containing 10% FBS (Invitrogen 10438026) and 1% GlutaMax (Gibco 35050-061). Cells were plated in a 24-well plate with a 30K density and induced to differentiate two days after initial plating with DMEM containing 10% FBS, Insulin (0.862 mM; Sigma, I2643) and Troglitazone (5 mM; Sigma, T2573). Cells were fixed with 4% PFA 5-7 days after initial differentiation for downstream analysis.

### Histology and immunofluorescence

Upon harvesting, TAs were processed as previously described (Johnson et al., 2022), where each TA was cut in two parts: for RNA isolation and histology. For histology, TAs were fixed in 4% PFA (paraformaldehyde) for 2.5hrs at 4°C, washed with PBS and cryopreserved overnight in a 30% sucrose solution. TAs were embedded in OCT-filled cryomolds (Sakura; 4566) and frozen in liquid N_2_-cooled isopentane. Sections were obtained with a Leica cryostat, collecting 3-4 sections that were 10µm thick every 250-350µm of the TA. For immunofluorescence, sections were blocked and incubated with primary antibodies in blocking solution (5% donkey solution in PBS with 0.3% Triton X-100) overnight at 4°C. Primary antibodies used were rabbit anti-Perilipin (1:1000; Cell Signaling, 9349 S), rabbit anti-MyoG (1:250; Proteintech Group 14688-1-AP), rabbit anti-cleaved Caspse 3 (1:500, Millipore Sigma AB3623), and goat anti-PDGFRα (1:250, R&D Systems #AF1062), rabbit anti-Ki67 (Abcam ab15580), chicken anti-GFP (1:1000, Avis lab). For the primary antibody rabbit anti-MYH3 (1:250, Proteintech 22287-1-AP), antigen retrieval was required, using a Sodium Citrate Buffer (10 mM Sodium Citrate, 0.05% Tween-20, pH 6.0). Secondary antibodies used were Alexa Fluor-conjugated from Life Technologies (1:1000), as well as directly conjugated dyes Phalloidin-Alexa 568 and 647 (1:200, Molecular Probes #A12380 & A22287), and incubated for 45min at room temperature. Slides were mounted (SouthernBiotech; 0100-01) before imaging. DAPI (Invitrogen, D1306) was used to visualize nuclei. To visualize fibrillar collagen, we performed the histological stain, Sirius Red. To detect lipid droplets in vitro, cells were stained with HCS LipidTOX Green Neutral Lipid Stain (Invitrogen H34475).

### Image Analysis

Images were acquired with a Leica DMi8 microscope equipped with a SPE confocal and high-resolution color camera. Images of the whole TA were obtained with the navigator function within the Leica LSA software. All images were quantified through ImageJ Software (v1.552p). To quantify percent of GFP^+^ adipocytes, total and proliferating FAPs, FAPs undergoing apoptosis and total MyoG cells, 6-7 random areas of one TA were imaged with a 20x objective and the total number of cells in the areas were averaged.

### Gene expression analysis

RNA was isolated as previously described (44). Briefly, muscle was homogenized using a bead beater (TissueLyser LT, Qiagen, 69980) in TRIzol (ThermoFisher Scientific, 15596026). To isolate RNA, chloroform was added, and RNA was purified following the QIAGEN RNeasy kit (74104) per manufacturer’s instructions. 500-800 ng of RNA was used to transcribe into cDNA using the qScript Reverse Transcriptase kit (Quanta bio; 84003). RT-qPCR was performed in technical triplicates or quadruplets on a QuantStudio 6 Flex Real-Time 384-well PCR System (Applied Biosystems, 4485694) using PowerUp SYBR Green Master Mix (ThermoFisher Scientific, A25742). Fold changes were normalized to the house keeping genes *Sra1* and *Pde12*.

### Cytokine Assay

After injury, TAs were harvested and lysed with a TissueLyser LT (Qiagen, 85600) as previously described, in 1x Phsphate buffered saline (PBS) with proteinase inhibitor (Pierce, Cat #A32953). Once centrifugated, the supernatant was collected, and protein concentration determined through a BCA assay (Pierce, Cat #232225). Samples were then run for an enzyme-linked immunosorbent assay (ELISA; 32-plex mouse cytokine assay, Eve technology).

### Statistics

The experimenter was blinded until data were collected. TAs with injury less than 50% of total area were excluded from this study. All data was graphed using GraphPad Prism with data presented as mean ± SEM. For comparing two samples with one variable, an unpaired two-tailed t test was used followed by a Holm-Šídák post hoc test if necessary. For more than two samples with one variable, a one-way ANOVA followed by a Dunnett’s multiple comparison test was used. For two variables, a two-way ANOVA followed by Tukey’s multiple comparison was carried out. A p value less than 0.05 was considered statistically significant and denoted as follows: *<0.05, **<0.01, ***<0.001 and ****<0.0001.

## Data Availability

The raw FastQ files are available on NIH GEO XXX. All other correspondence and material requests should be addressed to.

## Acknowledgements

The authors thank the members of the Kopinke laboratory for helping with data collection and critical reading of the manuscript. We also thank Karyn Esser for valuable input. This work was supported by the US National Institutes of Health (NIH) grants 1R01AR079449 to DK, 1R01HL171050 to DK and TER, T32HD043730 to AMN and F31HL174156 to VRP. DK was also supported by the Thomas Maren Junior Research Excellence Fund. The content is solely the responsibility of the authors and does not necessarily represent the official views of the NIH. All schematic figure models were created with BioRender.

## SUPPLEMENTAL FIGURES

**Supplemental Figure 1.**
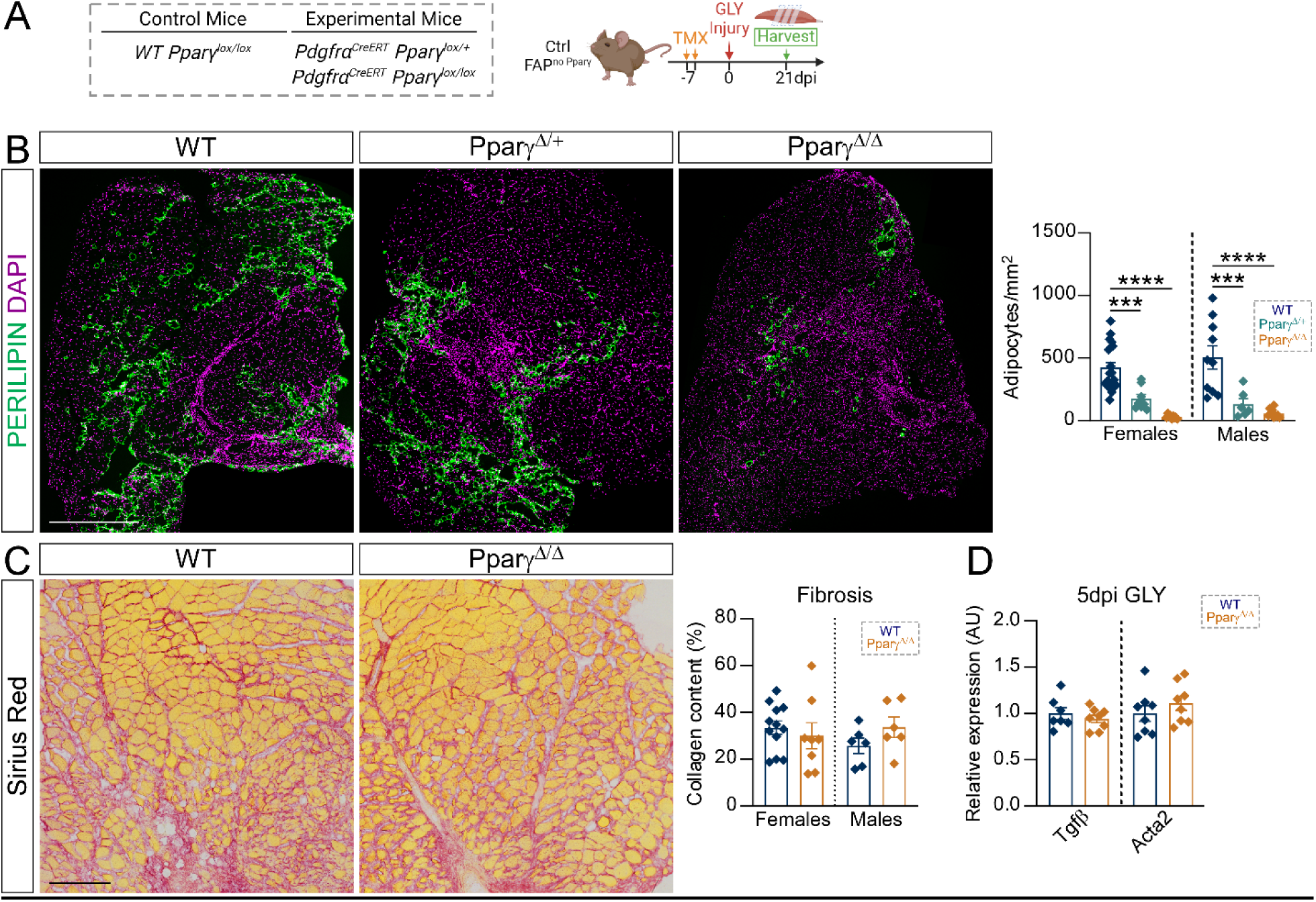
Pparγ is haploinsufficient but fails to impact injury-induced fibrosis. **A)** Experimental outline. **B)** Immunofluorescence of tibialis anterior (TA) cross-sections 21 days post GLY injury in WT, Pparγ^Δ/+^ and Pparγ^Δ/Δ^ mice stained for adipocytes (PERILIPIN, green). Nuclei were visualized through DAPI (magenta). Scale bar: 500µm. Quantification of adipocytes normalized to injured area 21 days after a GLY injury in females (WT n=20 TAs; Pparγ^Δ/+^ n=8 TAs; Pparγ^Δ/Δ^ n=9 TAs) and males (WT n=10 TAs; Pparγ^Δ/+^ n=6 TAs; Pparγ^Δ/Δ^ n=10 TAs). **C**) Differences in fibrosis was assessed by staining TA sections of WT and Pparγ^Δ/Δ^ mice for the histological stain Sirius red. Scale bar: 100µm. Quantification of the fibrotic index in females (WT n=12 TAs; Pparγ^Δ/Δ^ n=8 TAs) and males (WT n=6 TAs; Pparγ^Δ/Δ^ n=6 TAs). RT-qPCR of whole TA muscle lysate from WT (n=7-8 TAs) and FAP^no Pparγ^ (n=8 TAs) female mice 5 days post a GLY injury for the *Tgfβ* and *Acta2*. All data are represented as mean ± SEM. An unpaired two-tailed t test or a one-way ANOVA followed by a Dunnet’s multiple comparison was used.

**Supplemental Figure 2.**
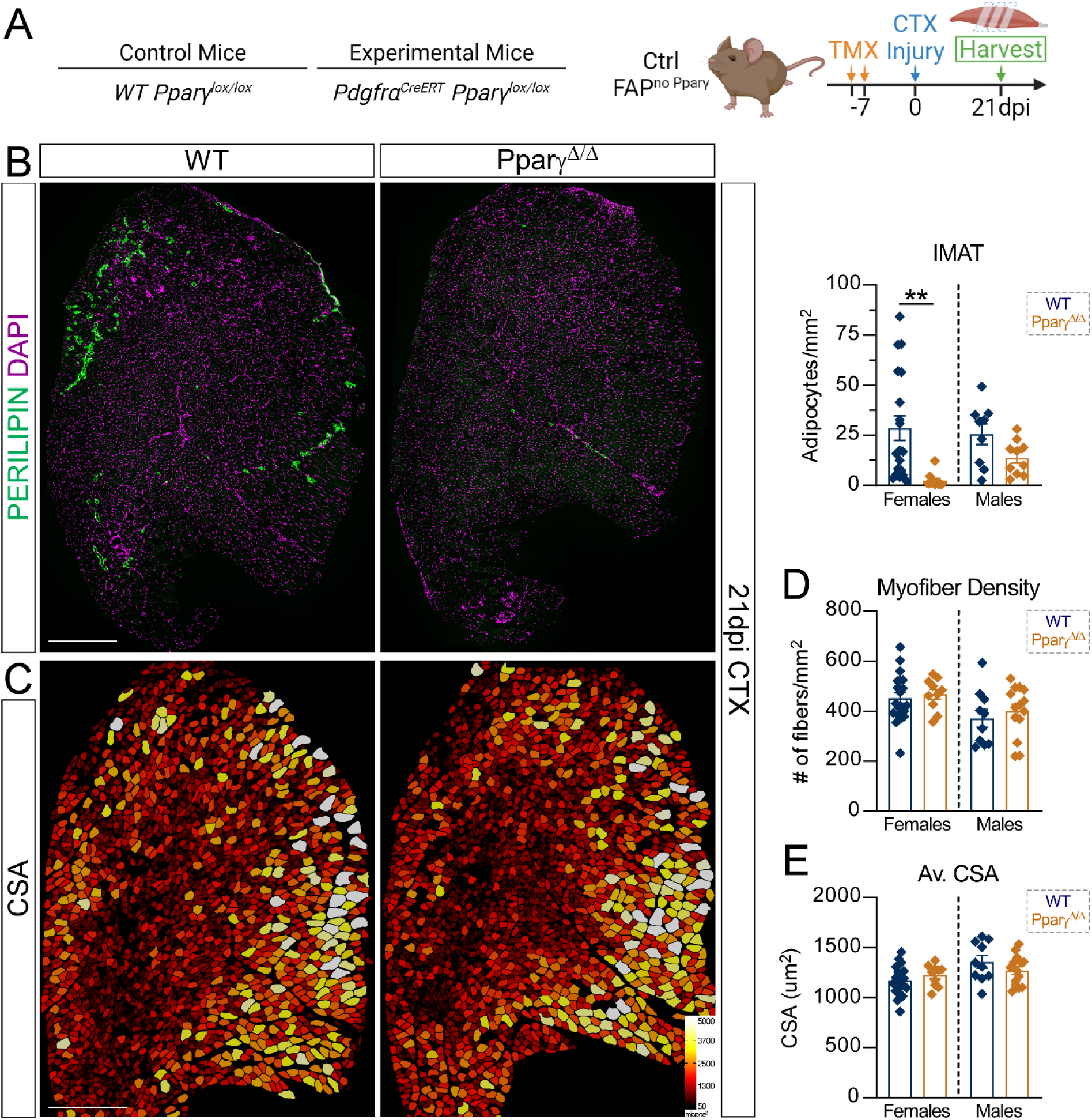
IMAT only inhibits muscle regeneration following a GLY but not CTX injury. **A**) Experimental design. **B)** Immunofluorescence of cross-section of tibialis anterior (TA) muscle from WT and FAP^no Pparγ^ mice 21 days after a CTX injury stained for adipocytes (PERILIPIN, green). Nuclei were visualized with DAPI (magenta). Scale bar: 500µm. Quantification of number of adipocytes normalized to injured area in females (WT n=18 TAs and FAP^no Pparγ^ n=10 TAs) and males (WT n=9 TAs and FAP^no Pparγ^ n=10 TAs). **C)** Myofibers were stained for PHAILLOIDIN followed by false color-coding according to size in WT and FAP^no Pparγ^ mice 21 days after a CTX injury. Scale bar: 500µm. **D**) Quantification of the number of fibers normalized to total TA cross-sectional area 21 days post CTX in females (WT n=20 TAs; FAP^no Pparγ^ n=10 TAs) and males (WT n=10 TAs; FAP^no Pparγ^ n=14 TAs). **E)** Quantification of average cross-sectional area (CSA) of muscle fibers in females (WT n=20 TAs; FAP^no Pparγ^ n=10 TAs) and males (WT n=10 TAs; FAP^no Pparγ^ n=14 TAs). All data are represented as mean ± SEM. A multiple unpaired two-tailed t test followed by a Holm-Šídák post hoc test was used.

**Supplemental Figure 3.**
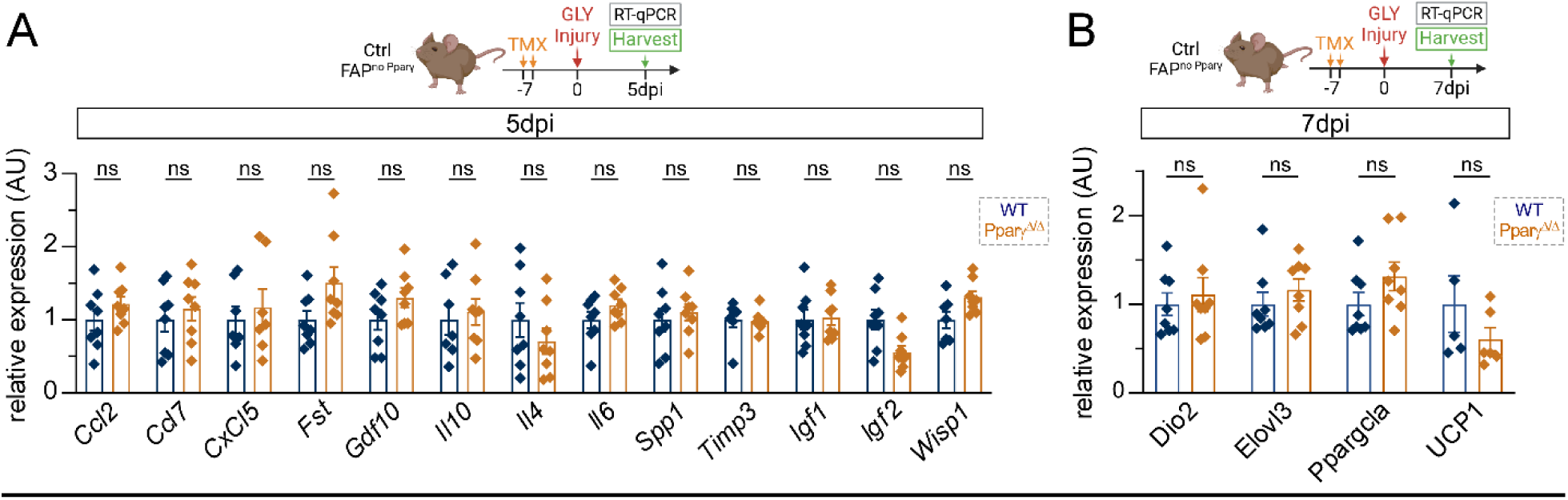
Removal of Pparγ from FAPs does not cause aberrant gene expression. **A)** Experimental outline. RT-qPCR of whole muscle TA lysate of female WT (n= 6-8 TAs) and FAP^no Pparγ^ (n= 6-8 TAs) mice 5 days post GLY injury for FAP-derived factors. **B)** Experimental outline. RT-qPCR of whole muscle TA lysate of female WT (n= 6-8 TAs) and FAP^no Pparγ^ (n= 6-8 TAs) mice 7 days post GLY injury for BAT-specific genes. All data are represented as mean ± SEM. A multiple unpaired two-tailed t test followed by a Holm-Šídák post hoc test was used.

